# Self-pressurized rapid freezing at arbitrary cryoprotectant concentrations

**DOI:** 10.1101/2022.12.30.522278

**Authors:** K. Rolle, K.A. Okotrub, I.V. Zaytseva, S.A. Babin, N.V. Surovtsev

## Abstract

Self-pressurized rapid freezing (SPRF) has been proposed as a simple alternative to traditional high pressure freezing (HPF) protocols for vitrification of biological samples in electron microscopy and cryopreservation applications. Both methods exploit the circumstance that the melting point of ice reaches a minimum when subjected to pressure of around 210 [MPa], however, in SPRF its precise quantity depends on sample properties and hence, is generally unknown. In particular, cryoprotective agents (CPAs) are expected to be a factor; though eschewed by many SPRF experiments, vitrification of larger samples notably cannot be envisaged without them. Thus, in this study, we address the question of how CPA concentration affects pressure inside sealed capillaries, and how to design SPRF experiments accordingly. By embedding a fiber-optic probe in samples and performing Raman spectroscopy after freezing, we first present a direct assessment of pressure buildup during SPRF, enabled by the large pressure sensitivity of the Raman shift of hexagonal ice. Choosing dimethyl sulfoxide (DMSO) as a model CPA, this approach allows us to demonstrate that average pressure drops to zero when DMSO concentrations of 15 wt % are exceeded. Since a trade-off between pressure and DMSO concentration represents an impasse with regards to vitrification of larger samples, we introduce a sample architecture with two chambers, separated by a partition that allows for equilibration of pressure but not DMSO concentrations. We show that pressure and concentration in the fiber-facing chamber can be tuned independently, and present differential scanning calorimetry (DSC) data supporting the improved vitrification performance of two-chamber designs.

## 1. INTRODUCTION

Suppression of ice-formation during cooling of biological samples for low-temperature preservation is typically accomplished by perfusing them with cryoprotectives agents (CPAs), solutes capable of lowering the freezing point of water and hence, the cooling rates required for achieving vitrification. However, because they tend to have a number of drawbacks, such as toxicity at high concentrations, alternative or complementary approaches have been developed. Among these, high pressure freezing (HPF)^[1]^ has found widespread adoption in electron microscopy, where suspending the sample in a CPA-rich medium can lead to undesirable structural distortions. HPF cools the sample by exposing it to a pressurized liquid nitrogen (LN) jet. At a target value of 210 [MPa], the melting point and homogeneous nucleation temperature of ice reach a minimum. Compared with freezing at ambient pressure, HPF affords two orders of magnitude lower critical cooling rates, which is said^[2]^ to translate to a 10-fold increase in permissible sample size. For pure water, one could hence expect vitrification to depths of ∼ 10 micrometers (albeit the HPF apparatus is generally^[3]^ not configured for this case), and for CPA-free biological samples to depths of < 300 micrometers (depending on cytoplasm viscosity).^[4]^

HPF has been a staple of electron microscopy labs for decades but requires a rather bulky and expensive machine, rendering it unsuitable in many other contexts, such as field work. As a simpler alternative, self-pressurized rapid freezing (SPRF)^[5]^ was introduced in 2009, where samples are placed inside sealed capillary tubes and pressure is generated when ice, whose density is lower than that of the liquid,^[6]^ forms synchronously upon plunging the capillary into a coolant (typically, LN). Studies have shown this to commence at the circumference defined by the capillary walls,^[7-8]^ leaving an ice-free region of interest in the capillary center where a sample can find refuge. While the relative size of the vitrified core has been proposed as a measure of the achieved pressure, these studies employed CPAs, and a halt to ice growth might accordingly only reflect the increased freeze-concentration of the solution. Thus while in HPF, applied pressure is reproducible and mainly depends on jet parameters, in SPRF, this quantity is generally unknown and studies typically limit themselves to empirically determining vitrification success.

Though CPA concentrations are often low in an electron microscopy context, the matter can be quite different if we examine SPRF with a view to reversible cryopreservation of live specimens. Indeed, the relevant studies^[9-10]^ have reflexively resorted to media with concentrations of 30 wt % or higher, and reported ambiguous results.^[11]^ This seems to confirm misgivings voiced in theoretical works^[12]^ to the effect that suppression of ice-formation by CPAs might have counterproductive consequences on pressure buildup inside the capillary. The matter has however not been conclusively investigated, foremost because of lack of direct access to the pressure value as mentioned, but possibly also because the case for HPF or SPRF of live samples might seem weaker at first glance. Indeed, requirements on sample quality here are generally less stringent than in electron microscopy, granting wider flexibility with regards to CPA choice and permissible CPA concentrations. More importantly, survival rates are often capped not by ice-formation during cooling, but rather during warming.^[13]^ The recent advent of pulsed laser warming techniques^[14-15]^ has however put much higher heating rates at the disposal of researchers. Hence, assumptions on minimum CPA concentrations^[14]^ and maximum sample sizes^[15]^ suitable for live cryopreservation have to be revisited, with the latter (high CPA, large sample) regime not yet catered to by existing SPRF container designs. Notably, SPRF capillaries could be an interesting substitute to the open cryocontainers (cryotop) from the cited experiments, since closed containers are sometimes preferred due to the lower risk of cross-contamination when several samples are stored in the same LN dewar.

In this paper, we hence propose to address the question of joint use of both CPAs and SPRF in the same samples, and how their cumulative effects can be harnessed to increase size limits on vitrification. To this end, we develop a method of direct pressure measurement inside sealed capillaries. For the latter, we follow SPRF standard practice in choosing copper tubes, which boast excellent thermal conductivity (and hence, cooling rates), high yield strength and ease of sealing by crimping. The drawback however is their lack of transparency, which impedes probing by optical methods, which would normally be particularly suitable due to their non-invasive character. As a solution, we embed an optical fiber inside the capillary; indeed, in process monitoring, fiber optic solutions are often advantageous due to their robustness, minimal space requirements and potential for remote readout. Point sensors for pressure are frequently realized by fabricating a Fabry-Perot cavity on the fiber endface.^[16]^ We will however adopt another solution and use Raman spectroscopy, which is a technique often available to cryopreservation labs for mapping various features of interest (CPA distribution,^[17]^ simple and eutectic^[18-19]^ crystallization, phase transitions,^[20]^ protein stability^[21]^ etc.) and observe the wavenumber shift of the hexagonal ice peak in the O-H band, which is well-known^[22]^ to exhibit high pressure sensitivity. This approach eliminates any need for sensor fabrication and will be seen to be particularly well adapted to the pressure range and sample compositions relevant in SPRF.

## 2. EXPERIMENT

Capillaries with dimensions capable of accommodating larger samples than in typical (low-CPA) SPRF (ID 0.86 [mm], OD 1.57 [mm]) and made from unannealed copper (#8117, K&S Precision Metals, Chicago, Illinois, U.S.A.) were chosen for the cryocontainer. They were chopped into 15 [mm] long segments by means of a capillary tube cutter (VALUE VRT-101), prized open by impaling on a 21G needle and cleaned by ultrasonicating in 0.2 wt % Tween 20. Then, they were individually strung onto < 0.5 [m] lengths of optical fiber whose endfaces had been prepared by a cleaver (Fujikura CT-11) and centered inside the capillary by threading through two telescoped 21G and 26G needles which protruded to a depth of 12 [mm] into it. The fibers were then retracted until > 3 [mm] stubs remain inside the tubes, and each was fixated by a drop of two-component glue (UHU Plus Endfest 300). Finally, the glue was left to harden overnight.

In view of the Raman experiment, multi-mode fibers (MMF) for power-delivery (105 [μm] silica core, 125 [μm] silica cladding, NA 0.22) were selected, where core dimensions are large enough for easy alignment and averaging over relatively large part of the sample chamber, but small enough not compromise signal at slit-widths that typically achieve adequate resolutions in dispersive Raman spectrometers (a limitation that can however potentially be overcome by slitless Fourier-transform Raman^[23]^ setups). Also in view of the Raman experiment, we chose dimethyl sulfoxide (DMSO) as a model CPA because it does not exhibit any overlap of its spectral features with those of water in the frequency range of interest (a problem commonly solved with other popular CPAs^[24]^ by using deuterated forms of either solute or solvent). The solution was prepared for various concentrations of DMSO (0-10 wt %) and poured into a shallow basin. The latter was slotted into a vertically mounted vice, fiberized capillaries were submerged in the basin and flushed with CPA solution that was injected into them via an L-shaped needle to expel air bubbles. Finally, they were sealed by positioning between a pedestal at the basin bottom and a piston that was pressed downwards by tightening the vice screw, removed from the basin, wiped dry and dropped vertically into LN.

In addition, slightly longer capillaries (18 [mm]) were prepared in analogous fashion, however, before submersion inside the basin, they were first flushed with a higher DMSO concentration (20 wt %) solution injected into them via a needle. Next, a piece of brass wire with ø 0.81 [mm] (#8160, K&S Precision Metals, Chicago, Illinois, U.S.A.) was inserted into the open end, a 2-3 [mm] long fragment was snipped off and then thrust to a depth of 8 [mm] into the tube center, so as to bisect it into two chambers of approximately equal volume (as a precaution against leakage, the wire was coated with a two-component PDMS glue, PK-68, T.U. 38.103508-81, after an adhesion promoter, P-11, T.U. 38.303-04-06-90, had been applied). Finally, residual high-CPA fluid was expelled from the open chamber by submersion in a basin with low DMSO concentrations (0-10 wt %) as above, capillaries were sealed, wiped dry and dropped immediately into LN to arrest diffusion.

The Raman experiment was conducted on a laboratory setup with 532 [nm] excitation wavelength at 250 [mW] power (Millennia II, Spectra-Physics, Santa Clara, CA), monochromator (SP2500i, Princeton Instruments, Trenton, NJ) with 500 [nm] grating pitch, NA=0.08 and LN-cooled CCD (Spec-10:256E/LN). At the spectrometer entrance, a f=200 [mm] lens focused scattered light onto a slit with 20 [μm] width, while a f=50 [mm] lens coupled the incident light into the fiber. For each sample, the accessible endface of the latter was cleaved and its position was adjusted on a 5-axis mount while maximizing the signal in the < 2000 [cm^-1^] spectral range, which is dominated by the fiber contribution. Then, the grating was turned so as to characterize the sample response in the 2000-4000 [cm^-1^] range. Afterwards, capillaries were rewarmed, opened with the capillary cutter and replunged into LN to obtain ‘non-pressurized’ reference spectra, while concentration could be verified from room-temperature data. Also, spectra of a neon-discharge lamp were acquired for wavelength calibration and of DCM dye for etalonning correction. The process was repeated until 5-10 spectra with pressure-evidence were available for each DMSO concentration, while cases bereft of any pressure shift (within the margin of error) were discarded from the statistics. A fresh sample was used for each pressurized spectrum.

Finally, reduced ice-formation in pressurized two-chamber capillaries with 20 wt % DMSO in the fiber-facing and 5 wt % DMSO in the other chamber was additionally confirmed from differential scanning calorimetry (DSC) traces acquired on a commercial calorimeter (DSC 200 F3, Netzsch). For this experiment, standard DSC pans were replaced with brass discs into both of which a groove matching capillary dimensions had been carved. Fiberized samples were prepared as for the Raman investigation and checked on the spectrometer. Those evincing successful pressure generation were sectioned in the middle with the capillary cutter that had been pre-cooled to LN temperature, then the fiber was trimmed and the CPA-rich half was inserted into the (also pre-cooled) brass disc. Finally, both capillary and mount were rapidly transferred together into the DSC chamber held at -160 [⁰C] and containing an empty but otherwise identical capillary half as reference. Under these conditions, the DSC scan was conducted at a heating rate of 5 [K/min]. For comparison, non-fiberized single-chamber capillaries with DMSO concentrations in the 20-40 wt % range were characterized also by DSC.

## 3. RESULTS AND DISCUSSION

Representative Raman spectra of quench-cooled samples templated on the traditional SPRF design^[5]^ and containing various DMSO concentrations are given in FIG. 1. Starting from the left (10 wt %) panel, the two sharp peaks at lower wavenumbers stem from the symmetric (2922 [cm^-1^]) and anti-symmetric (3009 [cm^-1^]) modes of CH_3_ stretching in DMSO,^[25]^ while the remaining portion of the spectrum represents ice. When comparing spectra of sealed (and hence, putatively pressurized) capillaries with those of open ones, the DMSO peaks are seen to remain immobile while the ice spectrum shifts to lower wavenumbers. At lower DMSO concentrations (middle plot), the relative intensity of ice peaks compared to DMSO is predictably larger. Indeed, when decreasing DMSO to 5 wt %, pressure is also seen to rise, as evidenced by the approximately twice larger wavenumber shift. While ice is known to form many exotic phases^[26-27]^ at low temperature, considering the target pressure of 210 [MPa] in SPRF, hexagonal (I_h_) ice is the most likely candidate although the low-density amorphous (LDA) ice spectrum looks very similar.^[28]^ Indeed, both reportedly^[29-30],[22]^ possess large pressure sensitivity, with dν/dP=159.8 [cm^-1^/GPa] for the hexagonal phase at -185 [⁰C]. The issue of non-standard ice does however become acute when decreasing concentration still further to 0 wt %. Here a minority of spectra (right panel) show a shift to longer wavenumbers, consistent with discontinuities in the pressure response^[22]^ reported upon phase transition at LN temperatures to crystalline (II, III, IX)^[30-31]^ and also (albeit less discontinuously so)^[32]^ LDA ice. While this would normally prevent a meaningful comparison with the standard case, the shown spectrum also comprises a smaller peak whose differential shift can be quantified at Δ(1/λ)=28.4 [cm^-1^] which is close to the maximum obtained from spectra with a more regular appearance. Nevertheless, the co-existence of two phases was not universal, so that all cases showing a jump to longer wavenumbers were excluded from the dataset.

**FIG. 1.**
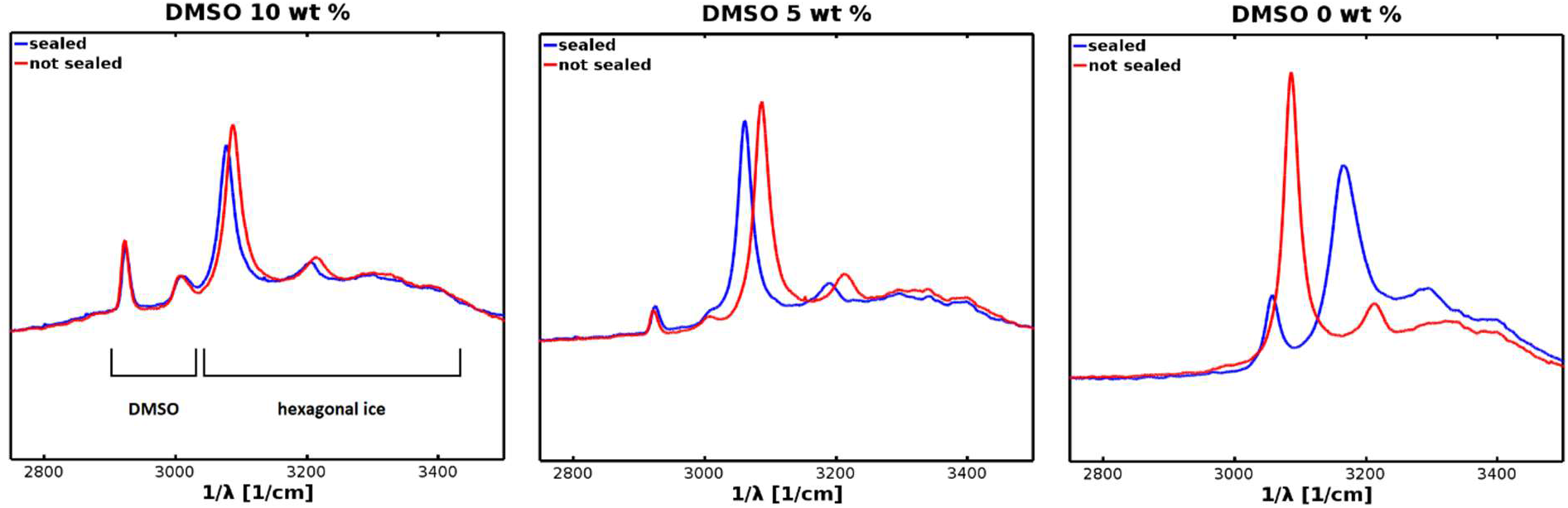
Raman spectra of aqueous DMSO solution in copper capillaries (ID 0.86 [mm], OD 1.57 [mm]) at liquid nitrogen (T=-196 [⁰C]) temperature, with DMSO concentration decreasing left to right. Blue spectra correspond to sealed capillaries, while red spectra were acquired after rewarming, cutting open and refreezing the same samples. The two left-most plots are representative while the right-most shows the atypical case (see discussion) of a two-phase system resulting from pressurization of pure water, with an increase in Raman shift in evidence for the dominant ice phase (by contrast, this mode is downshifted in the other two spectra).

Following this methodology, results for differential Raman shift and associated pressure as a function DMSO concentration are summarized in FIG. 2. Maximum values around 150 [MPa] are somewhat low considering the target (210 [MPa]), the yield strength of unannealed copper^[7],[33]^ and the confirmed^[34]^ operating range of the glue, possibly indicating that capillary wall thickness was too modest. Alternatively, uncertainty (by as much as a factor of two) on the Raman pressure derivatives recorded^[22],[30]^ in the literature could be a cause. Nevertheless, concerning the nonlinearity observed for the 0 wt % average value, failure of the seals is a more likely explanation than the just discussed exclusion of unassignable spectra from the dataset. However, average values for the remaining three concentrations show a linear decrease of pressure, with the fit intercepting the x-axis at 15 wt % regardless of second y-axis calibration. This value is not far from the limiting value in microscopy studies (20 wt % dextran)^[7]^ beyond which the (heterogeneously nucleated) ice shell along the inner copper wall was found to vanish. In contrast to the cited study, we seemingly fail to achieve vitrification at 15 wt % because of larger sample dimensions (see FIG. 5b below). In our case, suppressing heterogeneous freezing should thus not imply a much lower total ice fraction, but likely impacts crystal size distribution and location (along the metal surface), possibly important for producing the so-called^[7]^ “ice-jam” in the crimped end. Finally, another possible explanation for pressure drop-off is the CPA thermal contraction at higher concentrations^[6],[35-36]^ which could potentially nullify pressure built up in the earlier stages of freezing. However, assuming that the sample is fully frozen by homogeneously nucleated ice at 15 wt %, the maximally freeze-concentrated solution (60 wt % DMSO) should amount to no more than a quarter of weight. To fully compensate the thermal expansion of ice, its contraction would have to reach magnitudes of > 20 % volume change, while literature^[35]^ rather suggests a value of ∼ 10 % shrinkage at 60 wt %. Considering also that the 5-10 wt % concentrations from which we extrapolate are still lower, this effect is unlikely to significantly modify the observed 15 wt % limit. Thus, the pressure drop on exceeding it implies contribution from an additional mechanism (such as the ice-jam).

**FIG. 2.**
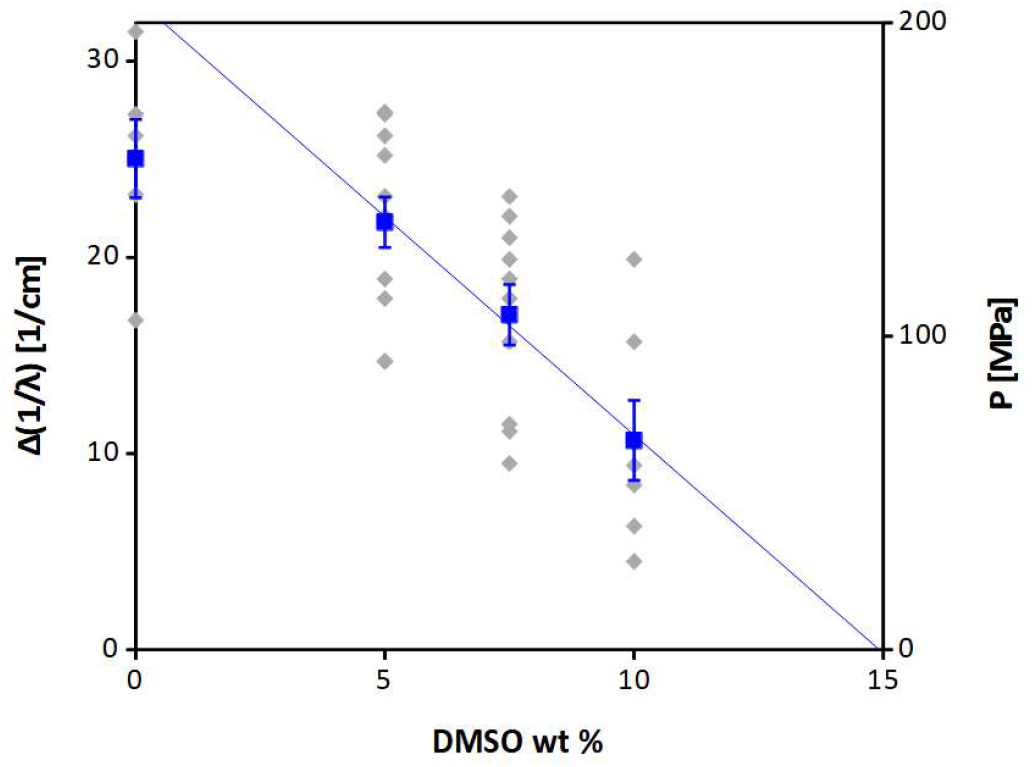
Differential Raman shift (left y-axis) and its translation to pressure (right y-axis, established from a tabulated^[30]^ value of dν/dP=159.8 [cm^-1^/GPa]) of the ice peak when comparing “sealed” and “unsealed” capillaries (FIG. 1) as a function of DMSO concentration. Grey diamonds represent individual spectra (see TAB. 1 for complete statistics), while blue squares are average values, and error bars the standard error of the mean. The average value representing pure water was excluded from the linear fit.

A consequence of this insight is that SPRF studies conducted at higher CPA concentrations in simple capillaries are likely doomed to fail. Hence, the remainder of our experiment focused on more sophisticated sample designs, as illustrated in FIG. 3 (left panel). Here, a wire fragment serves as a piston allowing ice formed in a low-CPA chamber to exert pressure upon a high-CPA one, all while preventing concentration equilibration between both. While in the related field of (slow) isochoric freezing the virtues of such a two-chamber approach to CPA distribution have already been investigated^[37]^ to the best of our knowledge this is the first SPRF implementation of this idea (though creative container formats have apparently^[38]^ been explored for other purposes). Representative spectra for the two-chamber design (FIG. 3) show a rather similar dependence of wavenumber shift on DMSO concentration (in the pressure-generating end) as before but differ in important aspects. First, relative intensity of the DMSO peaks is clearly stronger, as befits the CPA-rich (20 wt %) solution in the fiber-facing chamber. Secondly, pressurized and non-pressurized samples gave rise to rather similar relative intensities in the spectra of FIG. 1, whereas in FIG. 3 progression to the case of lower CPAs and higher pressure (5 wt %) seems to be accompanied by a reduction in the area of the ice peak (found to be by 32 % from a triple-Lorentzian fit, which was also close to the maximum achieved) albeit it could also just indicate less optimal quenching conditions in the reference case. Finally, the whole measurement is again repeated a statistically significant number of times for various DMSO concentrations, with results summarized in FIG. 4. Though somewhat less regular in appearance than FIG. 2, datapoints in the 5-10 wt % range are in almost identical positions, which is likely due to the low compressibility of water that prevents any damping even though the relative sample volume in which pressure is generated has been halved. Also, the absence of damping seems to suggest that CPA thermal contraction is not yet significant at the investigated concentration (similarly, the 2-chamber design allows for the ‘ice-jam’ hypothesis to be tested by switching around high and low concentrations, which will fail to produce any pressure). Moreover, no evidence of exotic ice phases was found for 0 wt % in the two-chamber spectra, confirming that the observed non-linearity in FIG. 2 is not just an artifact of data analysis.

**FIG. 3.**
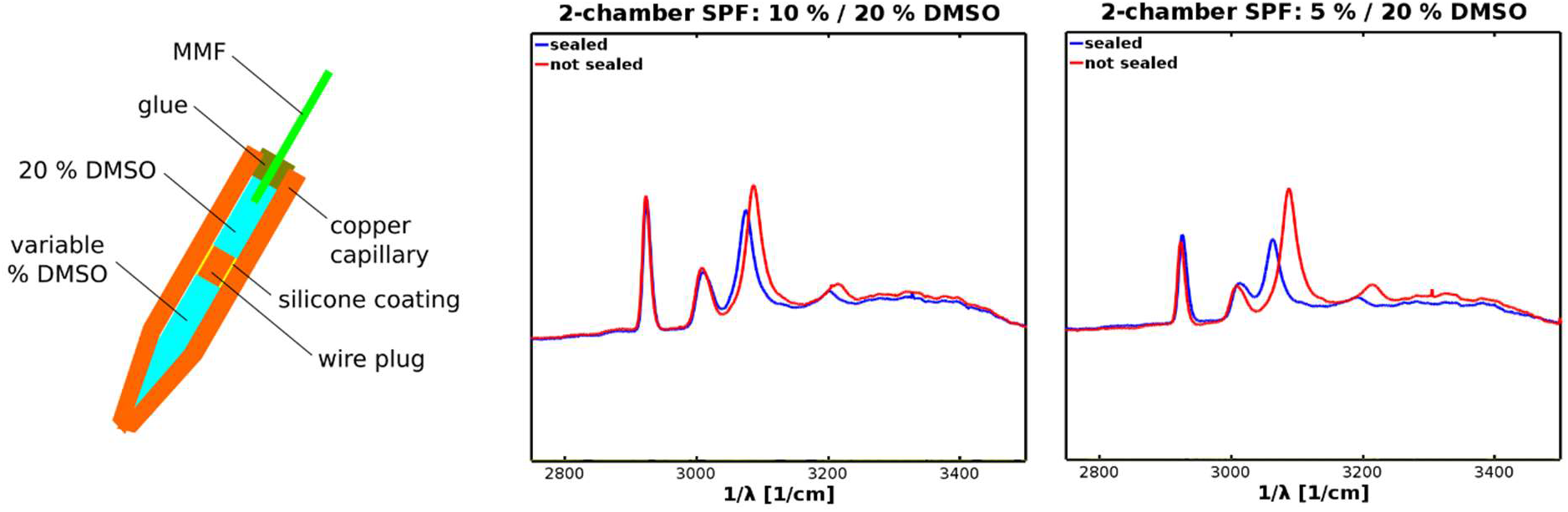
Double-chamber design (left) of self-pressurized copper capillaries with fiber-optic access and associated representative Raman spectra at liquid-nitrogen temperature (middle and right). DMSO concentration is constant (20 wt %) in the fiber-facing chamber of the capillary, and varies in the distal ends. Similarly to FIG. 1, blue spectra correspond to sealed capillaries, while red spectra were acquired after rewarming, cutting open and refreezing the same samples.

**FIG. 4.**
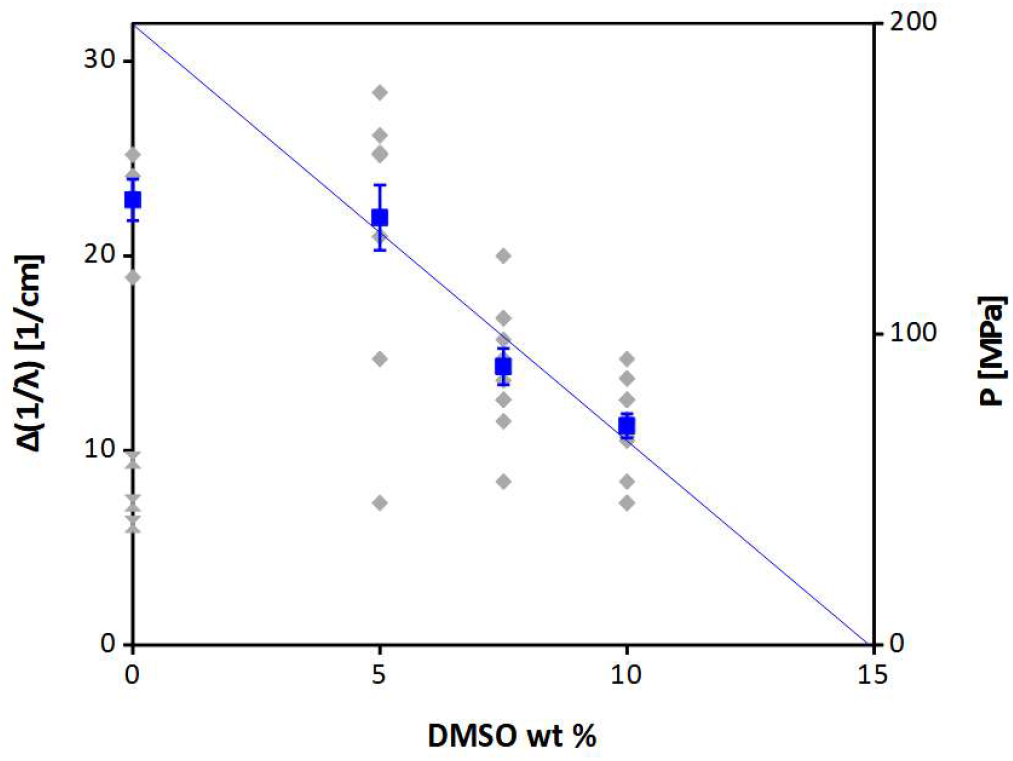
Similar to FIG. 2 but summarizing pressure dependence of a double-chamber design, as for samples and spectra under FIG. 3. For the case of pure water, the bottom 3 (hour-glass) datapoints were excluded from the corresponding average value.

Finally, pressure is not an end in itself when conducting SPRF, but meant to serve vitrification. Accordingly, we turn to DSC (FIG. 5 and TAB. 2) to check whether quench-cooled two-chamber capillaries will exhibit any recrystallization event upon warming. For the chosen capillary diameter, we can expect^[39-40]^ required DMSO concentration for vitrification^[41-42]^ to amount to at least 30 wt % (equivalent to an upper bound on the cooling rate < 1000 [K/s], however, the exact computation is not trivial^[12]^ because of its dependence on the ice fraction formed). The DSC trace for 20 wt % (FIG. 5a) exhibits a first, weak recrystallization event at a temperature T_ic_=-114 [⁰C] slightly above the glass transition (T_g_=-125 [⁰C]) and a much stronger one around -77 [⁰C]. The latter can be attributed^[43-44]^ to eutectic crystallization of the DMSO-water solution into a trihydrate crystal, which melts again at somewhat higher temperature (−65 [⁰C]). The eutectic crystallization event cannot in itself be taken as evidence of quench-cooling success because it supervenes even for the maximally freeze concentrated mixture (at 25 mol % or 60 wt %), albeit on a time-scale of at least several minutes.^[45]^ However, its absence from non-pressurized reference traces (FIG. 5b) hints at a greater quantity of liquid, unbound water and hence, faster diffusion during crystal formation. Less clear however is why said unbound water is apparently less prone to aggregating into ice when given the opportunity at lower temperatures, as in the reference traces. Possibly, the wire bolt continued to seal the opened end and prevented pressure relaxation above T_g_. The ambiguity is compounded by the only partially charted^[46-47]^ DMSO/water phase diagram at non-ambient pressure, as well as the rather short time-scales on which the DSC experiment can be conducted. However, the balance of evidence suggests that SPRF is indeed conducive to vitrification.

**FIG. 5.**
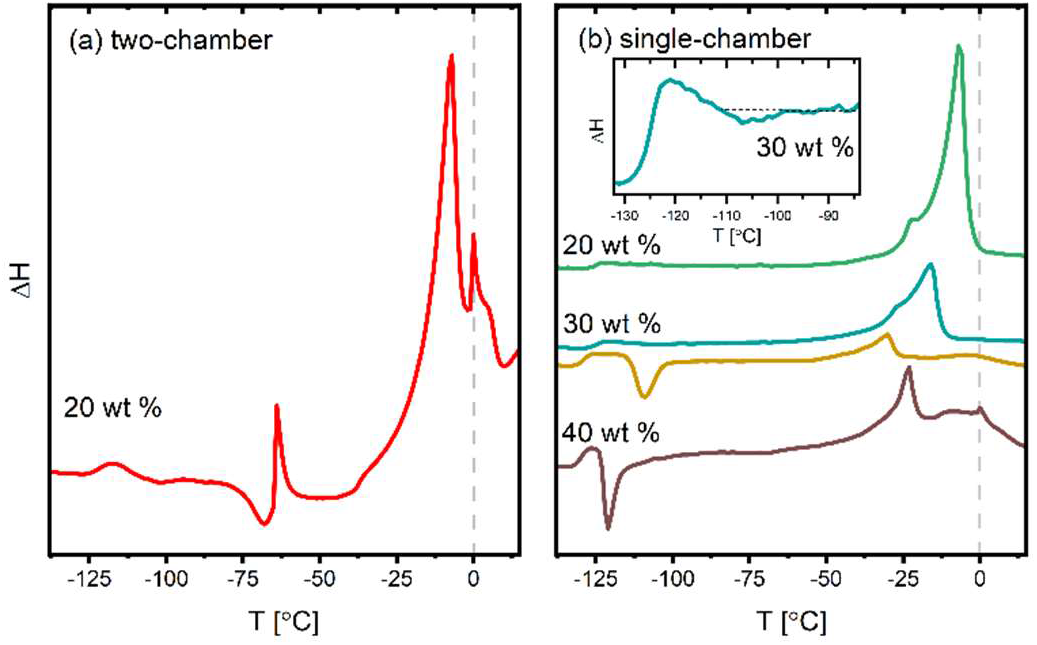
Differential scanning calorimetry (DSC) traces for quench-cooled samples with various DMSO concentrations, but juxtaposing a) two-chamber and b) single-chamber architecture (for the former, DMSO concentration was 5 wt % in the pressure-generating chamber, and verified wavenumber shift was 20 [cm^-1^]). Thermograms are vertically shifted for clarity. The inset in b) shows the rescaled weak feature in one of the DSC traces measured at 30 wt % DMSO, which is seen to be the limiting concentration below which no vitrification (barring for the maximally freeze-concentrated part) can be obtained without SPRF.

**TAB. 1.** Data from FIG. 2 (1-chamber design) and FIG. 4 (2-chamber design) with comprehensive statistical information on number of individual samples N (sample size), sum ∑, mean ∑/N and standard deviation σ (the error on the mean was computed as σ/√N). The 3 excluded spectra in 1-chamber 0 wt % case were found to have Δ(1/λ) < 0, as in FIG. 1 right, while in the 2-chamber 0 wt % case a subset of datapoints (indicated as hour-glass symbols in FIG. 4) was excluded since their values obviously placed them in a different ensemble.

| DMSO<br>wt % | 1-chamber |  |  |  | 2-chamber |  |  |  |
| --- | --- | --- | --- | --- | --- | --- | --- | --- |
| | N | $\sum$ [cm<sup>-1</sup>] | mean [cm<sup>-1</sup>] | $\sigma$ [cm<sup>-1</sup>] | N | $\sum$ [cm<sup>-1</sup>] | mean [cm<sup>-1</sup>] | $\sigma$ [cm<sup>-1</sup>] |
| 0 | 6 (+3) | 150.2 | 25.03 ± 2.00 | 4.89 | 5 (+3) | 114.4 | 22.88 ± 1.07 | 2.39 |
| 5 | 12 | 261.5 | 21.79 ± 1.29 | 4.46 | 12 | 263.6 | 21.97 ± 1.68 | 5.81 |
| 7.5 | 10 | 170.74 | 17.07 ± 1.54 | 4.89 | 11 | 157.4 | 14.31 ± 0.94 | 3.10 |
| 10 | 7 | 74.7 | 10.67 ± 2.04 | 5.39 | 14 | 157.7 | 11.26 ± 0.62 | 2.33 |

**TAB. 2.** Summary of DSC assays.

| Sample type | DMSO wt % | $T_g$ [°C] | $T_{ic}$ [°C] | $T_{ec}$ [°C] |
| --- | --- | --- | --- | --- |
| 2-chamber | 20 | -125±1 | -114±3 | -77±1 |
| 1-chamber | 20 | -128±1 | - | - |
| 1-chamber | 30 | -128±1 | -116±3 | - |
| 1-chamber | 30 | -132±1 | -115±1 | - |
| 1-chamber | 40 | -132±1 | -123.5±1 | - |

## 4. CONCLUSION AND OUTLOOK

The primary objective of this study has been to explore the cumulative benefits of combining SPRF and CPA, with a view to vitrifying larger sample volumes than each technique can handle separately. Our first important result does however concern SPRF with samples of all possible sizes. Indeed, it was experimentally demonstrated that naively increasing CPA content in order to improve vitrification will undercut pressure and any specific advantages SPRF derives from it, as has been suspected theoretically for some time.^[12]^ As per our second result, more sophisticated sample designs circumvent this trade-off, and can indeed be of relevance for vitrifying larger volumes in particular. The decrease in ice formation we found when doing so, as quantified by Raman spectra and corroborated by DSC, might seem rather modest but is largely in keeping with typical performance of SPRF or HPF (to the extent that comparison with previous SPRF studies is possible, which are largely microscopy-based). Nevertheless, in circumstances where other “CPA-free” freezing approaches e.g. based on open cryocontainers^[48-49]^ are off limits, the technical innovations introduced by our experiment could offer development opportunities worth exploring. On the one hand, the two-chamber design lends itself to a number of variations. E.g. wholly encapsulating one chamber using double-emulsion approaches^[50]^ could make chambers less prone to interdiffusion or (if sample volumes are minimized) damping, and their construction less labor-intensive. Alternatively, the cell walls in the sample itself could define a chamber, if either ice-nucleating agents are deployed externally to tune pressure^[51]^ or non-penetrating CPAs internally (by means of typical^[52]^ drug-delivery techniques). On the other hand, pressure monitoring through a fiber-optic Raman probe is the second important innovation. Due to almost negligible material cost, it could be suitable for routine deployment in a variety of related process monitoring contexts e.g. isochoric freezing of pharmaceuticals in vials;^[53-54]^ for the immediate needs of SPRF, it should allow for implementation of post-selection strategies to diminish the large statistical variations we witnessed (FIG. 2 and FIG. 4). Finally, fiber-optic monitoring could also become entrenched if the fiber can fulfill a double-role for other purposes. Notably it could facilitate power delivery during pulsed laser warming^[15]^ which would relax constraints on laser beam quality, an issue with widely available diode sources. Indeed, since rates from convective heating alone are incompatible with live recovery of large samples, any protocol seeking to expand vitrification volumes from quench-cooling, such as the one we just developed, should benefit from ease of integration with pulsed laser warming techniques.

## ACKNOWLEDGMENTS

This work was supported by Russian Science Foundation (Grant No. 21-72-30024). Part of the experiments was performed in the multiple-access center “High resolution spectroscopy of gases and condensed matters” in IA&E SB RAS (Novosibirsk, Russia).

